# Investigation of DPPC liposomes reveals their capability to entrap Aroclor 1260, an emerging environmental pollutant

**DOI:** 10.1101/829218

**Authors:** Andrew Lozano, Monica D. Rieth

## Abstract

Persistent organic pollutants (POPs) are a class of organic compounds that can accumulate in biological and ecological environments due to their resistive nature to chemical, thermal and photo degradation. Polychlorinated biphenyls (PCBs) are a class of man-made POPs that saw wide-spread use in commercial and industrial infrastructure as both an insulator and coolant in electrical transformers and capacitors. 2,2’,3,3’,4,4’-hexachlorobiphenyl (HCBP) was one of the most widely produced PCBs. As these mechanical structures fail or are decommissioned, PCBs are released into the soil, migrate to the water table, and eventually spread to nearby ecosystems by rain and wind. The stability of POPs and specifically PCBs leave few options for environmental waste removal. Conventionally, liposomes have been used for their drug delivery capabilities, but here we have chosen to investigate their capability in removing this class of emerging environmental pollutants. Liposomes are small, nonpolar lipid bi-layered aggregates capable of capturing a wide variety of both polar and nonpolar compounds. Dipalmitoylphosphatidylcholine (DPPC) is a well-characterized lipid that can be derived from natural sources. It is a phospholipid typically found as a major component of pulmonary surfactant mixtures. Liposomes were prepared using probe-tip sonication for both direct and passive incorporation of the HCBP compound. Assimilation was assessed using both differential scanning calorimetry and UV-Vis spectroscopy. After direct incorporation of HCBP the phase transition temperature, T_m_, decreased from 40.8 °C to 37.4 °C. A subsequent UV-Vis analysis of HCBP by both direct and passive incorporation showed an increase in HCBP incorporation proportionate to the length of exposure time up to 24 hours and relative to the initial quantity present during the direct incorporation. Together the decrease in T_m_ and increase in absorbance are indicative of HCBP incorporation and further demonstrate the potential for their use as a method of sustainable environmental cleanup.

## Introduction

Liposomes are spherical-shaped vesicular nanoparticles that have tremendous potential in biomedical and bioengineering applications. They are bilayered nanostructures often comprised of phospholipids that form an aqueous core where small molecules can be encapsulated. Lipids can vary in both their chemical and physical properties giving rise to larger nanostructures with unique properties unto themselves. Lipids with varying chain lengths and different degrees of saturation can be introduced to tune them for different applications.^1–5^ They are readily prepared using techniques like extrusion, sonication and rapid ethanol injection. Each of these methods influences properties such as size (diameter), lamellarity (single bilayer or multi-bilayer), and polydispersity (range of sizes).^6–8^ Once prepared their phase behavior, drug permeability, and thermal stability can be investigated and characterized.^9,10^ Other physical properties of interest include surface charge (zeta-potential) and bilayer fluidity.^11–13^ They can also be used in molecular biology to facilitate organism transformation and transfection with foreign DNA or RNA.^14–17^

While they continue to be extensively studied for their applications in biomedical and bioengineering less is known about their use for environmental purposes. Here we used liposomes prepared with 1,2-dipalmitoyl-*sn*-glycero-3-phosphocholine, DPPC, to introduce a polychlorinated biphenyl compound in order to assess the capacity for these liposomes to entrap environmental pollutants. DPPC has reportedly been used to prepare liposomes for drug delivery and has also been investigated for its effects on membrane stability and permeability in biophysical studies. Other compounds like cholesterol, have also been incorporated which stabilizes liposomes under certain conditions. These effects can vary depending on other factors like liposome chain length and relative cholesterol abundance.^12,18–23^ Interestingly, even small changes to conditions like preparation, pH, lipid chain-length and heterogeneity are sufficient to alter the physical behavior of liposomes and affect important aspects of their drug permeability and controlled release.^10,24,25^

Here we investigated the use of liposomes as a vehicle for the absorption of compounds posing a potential environmental hazard. One class of compounds considered to be a growing concern are persistent organic pollutants (POPs).^26–29^ These compounds can slowly leach into the ground water where they are deposited into the soil eventually leading to contamination of surrounding water sources. A major subclass of persistent organic pollutants are the polychlorinated biphenyl compounds, PCBs, which share a similar basic structural motif containing chlorine atoms substituted at various positions (2-10) about a biphenyl ring (Figure 1). These compounds are remarkably stable and resistant to environmental degradation causing them to accumulate and pose serious environmental and health concerns. They have been used industrially for their desirable electrical insulating and heat transfer properties as well as plasticizers in paint and other polymer-based commercial products. They are also reportedly released into the ground from landfill waste sources such as microplastics and can be released into the air upon incineration of waste materials posing respiratory dangers as well (Figure 2).^30–32^ The origins of PCBs in the environment extend beyond industrial sources and became a growing concern in the early 1990s when they were discovered in commercial paint pigments.^33–36^ Only now are we becoming increasingly aware of their potential threat to the environment and human health. Once exposed, PCBs, also known by their commercial name, Aroclor, can cross the cell membrane and bind with receptors in both human and mouse models leaving organisms susceptible to its unpredictable and sometimes negative effects.^37–42^ They can also have significant environmental impacts by altering the local ecosystem and are believed to promote the growth and invasiveness of microbial species like cyanobacteria leading to formation of algal blooms.^43^

**Figure 1.**
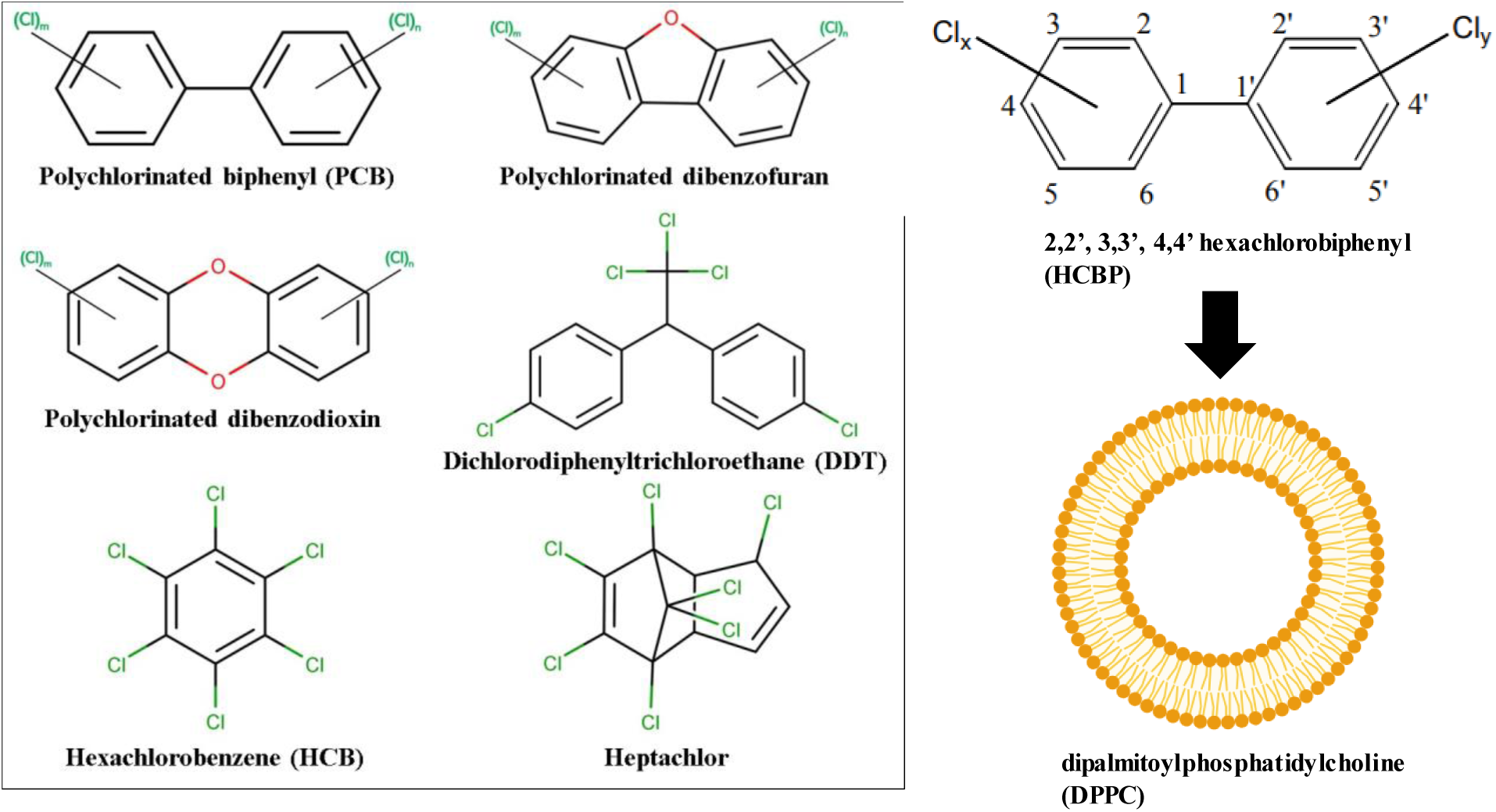
Common classes of persistent organic pollutants. General structure of a polychlorinated biphenyl (PCB) compound. Aroclor 1260 is substituted at positions 2,2’, 3,3’, 4,4’ to give a hexachlorinated biphenyl compound (HCBP). DPPC forms a bilayered lipid structure that can capture small molecular compounds.

**Figure 2.**
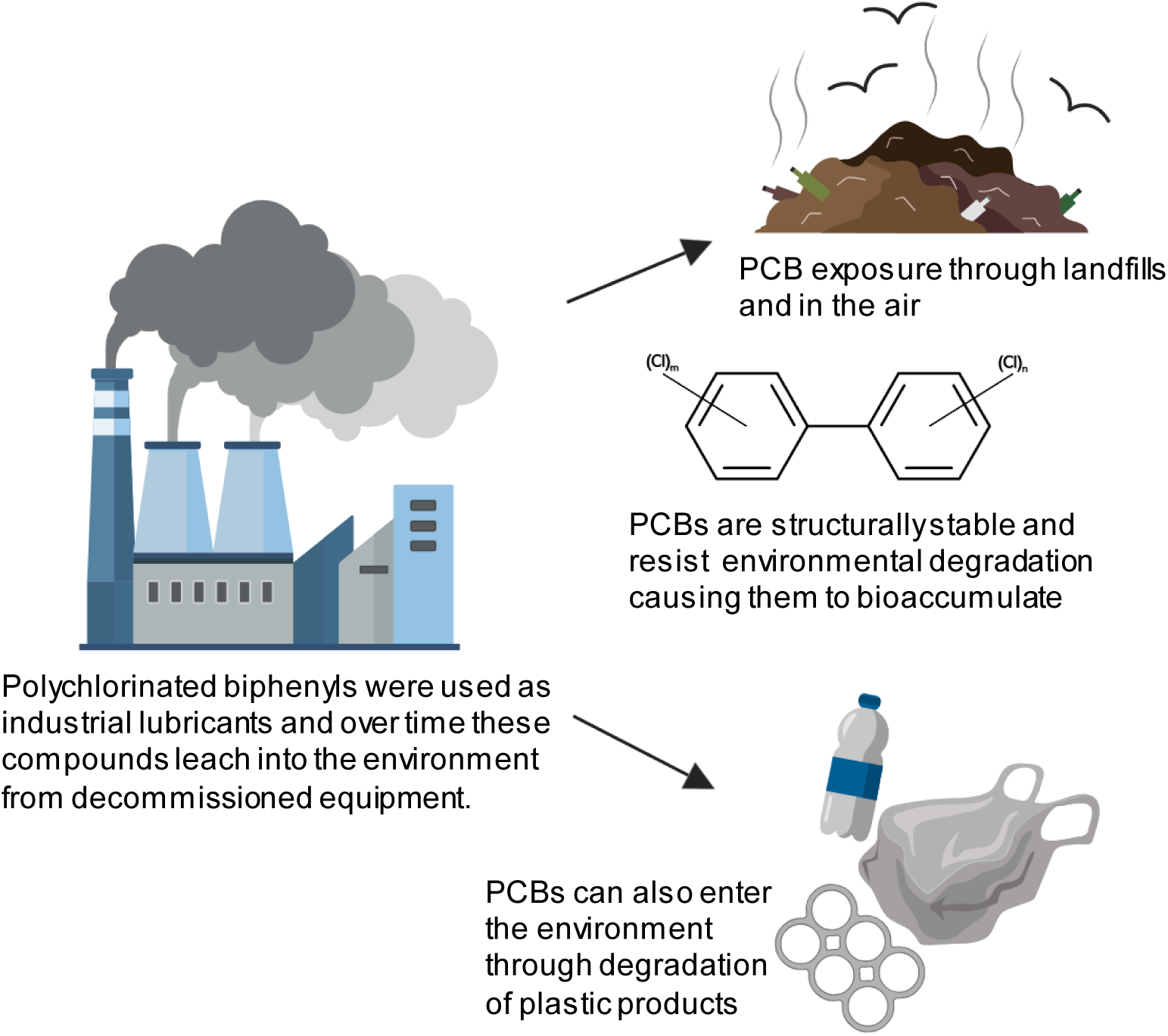
Schematic diagram representing the generation and life cycle of many PCBs. Their predicted bioaccumulation poses potential risks to the health of the surrounding ecosystem.

Current technologies have been adapted to address this issue, but here we report one of the first instances using a biomaterial-based approach to capture these compounds. We sought to capture a polychlorinated biphenyl compound, Aroclor 1260, also known as 2,2’,3,3’, 4,4’-hexachlorobiphenyl (HCBP), using pure DPPC liposomes. This compound is a congener of the polychlorinated biphenyl family and can induce human receptor activation and mimic the role of adipose tissue in hormone signaling and reproductive processes as well as patterns of protein expression.^38,41,44^ We monitored this process by measuring changes in the thermal stability of the resulting liposome mixtures in conjunction with a spectrophotometric analysis to track HCBP.

Differential scanning calorimetry is a powerful biophysical method that can be used to assess the stability and extract thermodynamic properties of protein-protein interactions, protein-lipid interactions, lipid-lipid interactions, protein-nucleic acid interactions and carbohydrate-lipid interactions. It can also be used to monitor protein unfolding and gain insight into factors that stabilize protein structure.^17,46,47^ Using this approach we found that increasing concentrations of HCBP in our liposome preparations generally broadened and lowered the characteristic transition temperature, T_m_, previously reported for pure DPPC liposomes. Further, a UV-Vis spectrophotometric analysis revealed that this compound readily incorporates into liposomes in both a direct manner when they are co-dissolved and prepared together and also passively when pre-formed liposomes are exposed to the compound. This system may be useful for the pretreatment of wastewater and potable water where current methods are unable to extract these types of compounds.^48,49^

## Materials and Methods

### Preparation of DPPC Liposomes with 2,2’, 3,3’, 4,4’-HCBP

The saturated lipid, 1,2-dipalmitoyl-sn-glycero-3-phosphocholine (DPPC), was purchased from Avanti Polar Lipids (cat.# 850355, Alabaster, AL). No further purification and characterization was necessary. Ampules containing 2,2’,3,3’,4,4’-hexachlorobiphenyl solution (Aroclor 1260) at a concentration of 1 mg/mL dissolved in hexanes were purchased from AccuStandard^®^ (cat# C-260S-H-10X, New Haven, CT) The compound was used without further purification.

DPPC liposomes were prepared at a total lipid composition of 25 mg/mL. Mixtures were prepared based on the following molar ratios of HCBP to DPPC: 0, 1, 5, 10, 20%. To prepare samples, DPPC was weighed out using an analytical balance to a mass of 25 ± 0.2 mg. The calculated mass of dry DPPC was weighed into a 2 mL glass screw top vial followed by the addition of 1 mg/mL Aroclor 1260 solution. The vials were then back-filled to a total volume of 1 mL with 200-proof ethanol. Each sample was prepared according to the calculated ratios of HCBP:DPPC and summarized in Table 1. The dried lipid films were stored long-term at −20 °C.

**Table 1.** Preparation of Molar Ratios of DPPC and 2,2’, 3,3’, 4,4’-hexachlorobiphenyl (HCBP)

| **Molar Ratio**<br>**HCBP/DPPC**<br>**(mol%)** | **Mass of**<br>**DPPC (mg)** | **Concentration**<br>**of DPPC**<br>**(mM)\*** | **Total volume**<br>**of HCBP (μL)** | **HCBP (mg)** | **Total volume**<br>**of EtOH (μL)** |
| --- | --- | --- | --- | --- | --- |
| **0** | 25.0 | 34.1 | 0.00 | 0.00 | 1000 |
| **1** | 24.9 | 33.9 | 122.9 | 0.123 | 877.1 |
| **5** | 24.4 | 32.8 | 614.5 | 0.615 | 385.5 |
| **10** | 23.8 | 32.0 | 1229.0 | 1.229 | 385.5 |
| **20** | 22.5 | 30.3 | 2458.2 | 2.458 | 180.6 |

**Table 2.**
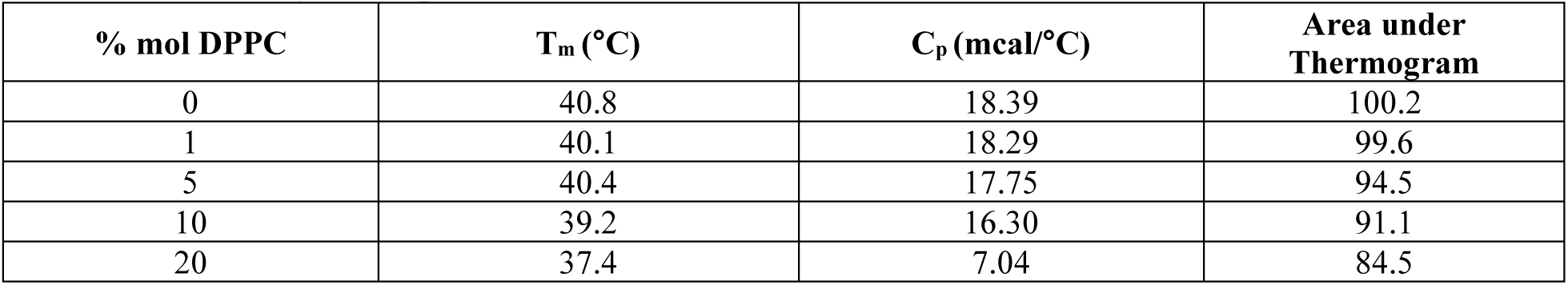
Thermodynamic parameters from DSC.

| % mol DPPC | $T_m$ (°C) | $C_p$ (mcal/°C) | Area under Thermogram |
| --- | --- | --- | --- |
| 0 | 40.8 | 18.39 | 100.2 |
| 1 | 40.1 | 18.29 | 99.6 |
| 5 | 40.4 | 17.75 | 94.5 |
| 10 | 39.2 | 16.30 | 91.1 |
| 20 | 37.4 | 7.04 | 84.5 |

To prepare liposomes from the dried lipid films, 1.0 mL of 20 mM HEPES, 100 mM NaCl pH 7.4 was added to the glass vial containing the lipid film. Samples were vortexed to mix and hydrate the lipid-HCBP film. This resulted in a milky white suspension that contained some larger white particulates. To suspend the lipids more homogeneously the mixture was sonicated for 2 minutes using a probe-tip sonicator (Fisher Scientific, Hampton, NH) set to 20% duty cycle with a pulse time of 2 seconds followed by a rest period of 2 seconds. One cycle was sufficient to homogeneously suspend the lipids to a milky white liquid with no visible large white particulates. The cycle was repeated three additional times. A total of four cycles at 2 minutes per cycle was carried out on each sample (8 minutes total). A 2-second rest period between pulses was incorporated to prevent excessive heating of the lipid mixtures. Cooling the liposomes was kept to a minimum to avoid dropping too far below the T_m_ of DPPC, 41.0 °C, which can inhibit liposome formation.

The supernatant was transferred to a clean 2.0 mL Eppendorf tube and centrifuged for 3 minutes at 10,000 rpm. The supernatant was removed and again transferred to a clean 2.0 mL Eppendorf tube. Samples were stored overnight at 4 °C and DSC studies were carried out the following day. Samples were not prepared more than 16-24 hours in advance of the DSC studies to preserve sample integrity and minimize liposome degradation.

### Differential Scanning Calorimetry (DSC) scanning parameters

Measurements were carried out on a VP-DSC high sensitivity scanning calorimeter (MicroCal, Northampton, MA, USA). All samples were scanned at a rate of 60 °C/hr beginning at 20 °C and ending at 70 °C. Samples were pre-equilibrated for 5-10 minutes at 20 °C (approximately room temperature) prior to the initial scan. The raw data were saved and plotted using KaleidaGraph version 4.5 scientific graphing program (Synergy software, Reading, PA). Prior to DSC analysis, stored liposome samples were removed from the refrigerator and left to equilibrate at room temperature for at least one hour. The samples were centrifuged at 10,000 rpm for 3 minutes to remove any unincorporated lipids. The supernatant was carefully transferred to a clean 2.0 mL Eppendorf tube. All samples were degassed for approximately 30 minutes along with 20 mM HEPES, 100 mM NaCl pH 7.4 buffer. HEPES buffer was chosen due to its pH stability over a broad temperature range. A scan of the buffer was acquired and collected and used for baseline subtraction. Due to concerns with irreversible degradation, one scan per sample was obtained and sample replicates were carried out on freshly prepared samples following the procedure outlined above.

### Preparation of passive diffusion of DPPC liposomes with 2,2’,3,3’,4,4’-HCBP

Four samples of 25 mg/mL DPPC were prepared separately from HCPB by weighing 25 mg of dry DPPC powder into a 2 mL glass screw top vial followed by the addition of 1.0 mL of a 20 mM HEPES, 100 mM NaCl pH 7.4 buffer. Liposomes were then prepared from the samples using the probe-tip sonication method outlined above. Four samples and one control sample of 1 mg HCBP were prepared by adding 1.0 mL of HCBP solution (1 mg /mL in hexanes) into a 2 mL screw top glass vial and drying down under a steady stream of nitrogen gas until a dry film of HCBP appeared and constant mass was achieved. The previously prepared 25 mg/mL DPPC liposomes were then added to four of the 2 mL glass vials containing the dried HCBP film and placed on an end-over-end mixer for 4, 8, 12, and 24 hours to allow thorough exposure of the liposomes to the dried HCBP film. The control sample was combined with 1.0 mL of 20 mM HEPES, 100 mM NaCl pH 7.4 buffer only, and placed on the end-over-end mixer for 24 hours. At each time interval, the samples were recovered from each vial and transferred to a 2 mL Eppendorf tube for analysis by DSC using methods described above. Samples were also retained for UV-Vis analysis.

### UV-Vis spectroscopy analysis of 2,2’, 3,3’, 4,4’-HCBP

The UV-Vis absorbance of 2,2’, 3,3’, 4,4’-HCBP was assessed using a Cary 300 UV-Vis spectrophotometer (Agilent Technologies, Santa Clara, CA). The maximum absorbance was measured using an HCBP sample prepared by transferring 1.0 mL of the 1 mg/mL stock HCBP hexane solution to a 2 mL glass vial and drying down under a steady stream of nitrogen gas until constant mass was achieved. To the glass vial was added 1.0 mL of 200 proof ethanol and the contents were mixed for 1 minute using a benchtop vortex mixer. In a quartz cuvette, 200 proof ethanol was used for both reference and sample cells and scanned from 800 nm to 200 nm. Lambda max was found based on the maximum absorbance and corresponding wavelength.

Next, a standard curve was generated and the extinction coefficient was determined using the standard Beer-Lambert relationship. To prepare samples for the standard curve analysis solutions of 20, 15, 10, 8, 6, 4, 2, and 0.5 μg/mL HCBP in ethanol (200 proof) were prepared from a working stock solution of 100 μg/mL. The eight samples were measured at a wavelength of 236 nm using a standard benchtop UV/Vis spectrophotometer. Data were processed and the extinction coefficient was determined from the slope of the standard curve.

To measure HCBP directly incorporated into liposomes, samples from the DSC were recovered. To a semi-micro quartz cuvette 780-790 μL of 200-proof ethanol was added followed by 20-10 μL of the sample recovered from the DSC analysis. The sample was mixed well to solubilize the liposomes and the absorbance was measured at 236 nm. Absorbance measurements were kept between 0.2 and 1.2 and sample dilution were made accordingly.

Samples from the passive incorporation were recovered after 4, 8, 12 and 24-hour time points and diluted into 780 μL 200 proof ethanol in a quartz cuvette to a final volume of 800 μL. Absorbances were measured at 236 nm and % incorporation was determined from the 1 mg dried film. Residual HCBP was recovered from the inside of each vial by flushing each vial twice with 20 HEPES, 100 mM NaCl pH 7.4 buffer. Ethanol was added to dissolve contents and 3 μL was diluted in a quartz cuvette and the absorbance was measured. From the absorbance measurement and the calculated extinction coefficient the concentration of HCBP was found and % remaining could be determined.

## Results

After each liposome preparation samples were centrifuged to remove unincorporated lipids and small bits of titanium from the sonicator probe. After centrifuging, a white pellet was visible at the bottom of the microfuge tube, which became more readily apparent in samples that contained higher amounts of HCBP. Samples were stored overnight at 4 °C to preserve sample integrity until measurements could be carried out, but not longer than 24 hours. Liposomes are only stable for a relatively short period of time before they begin to degrade and constituent lipids begin to precipitate out of solution.^18,20,50^ After samples were removed from 4 °C and left to equilibrate at room temperature, they were centrifuged again at 10,000 x g for 3 minutes and a minimal white pellet was visible in all samples. Most of the samples were prepared ahead of time and stored overnight. DSC scans were carried out beginning at room temperature to avoid exposing the lipids to temperatures far below the T_m_ for DPPC, since cooler temperatures can also affect the fluidity of the lipid tails and accelerate liposome degradation.^51^ The initial quantity of DPPC used for these experiments was previously optimized to ensure that an appreciable signal arising from the T_m_ would be captured. We found that 10 mg/mL and 25 mg/mL both gave the best signal, and subsequent studies were carried out using 25 mg/mL DPPC to also increase the loading capacity of the compound.

In Figure 3, panels A and B a small peak was visible in the DSC thermograms, which we attributed to residual unincorporated lipid that was not completely removed during centrifugation. In previous experiments samples prepared with 2, 5, 10 mg/mL DPPC following the method of sonication showed a noticeable decrease in this small peak with DSC analysis, which suggests what we believe to be unincorporated lipid varies in proportion to the total amount of lipid in the sample (manuscript under review). Filtering the sample was avoided to minimize the risk of disrupting and altering the physical properties of the liposomes.

**Figure 3.**
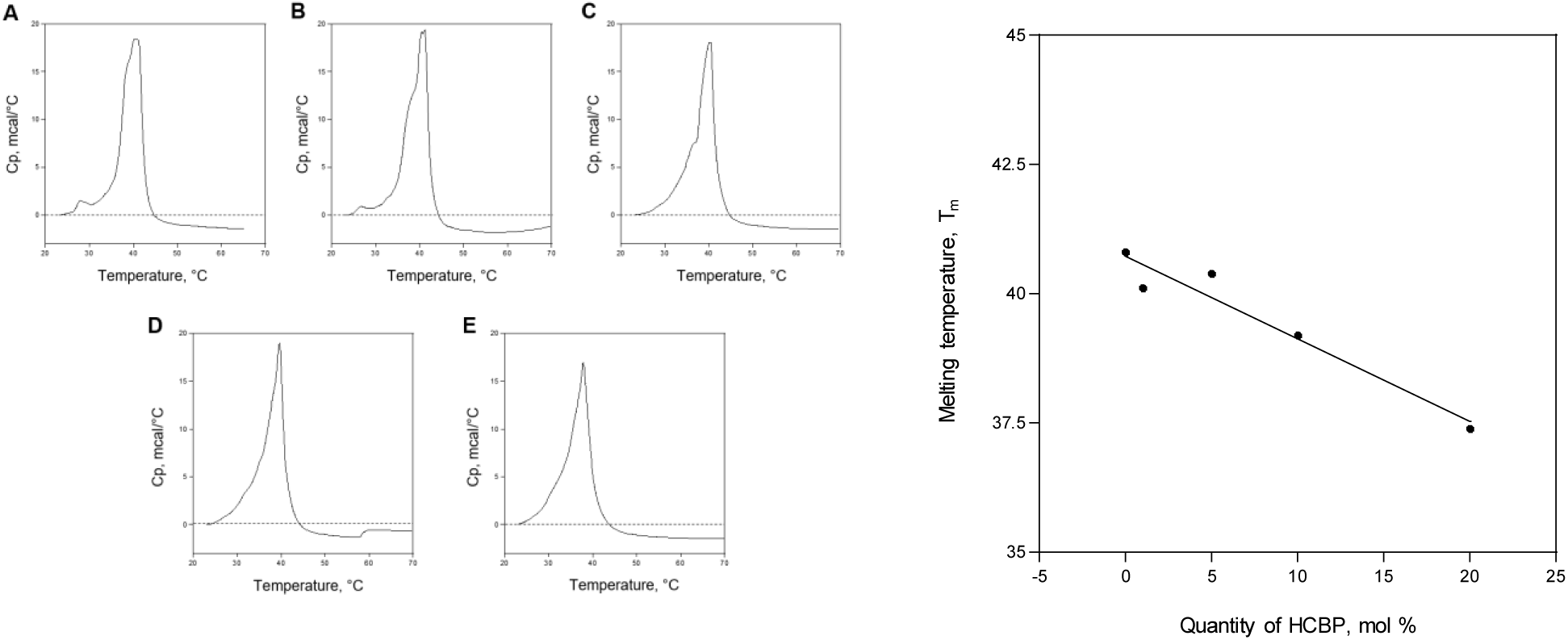
Differential scanning calorimetry (DSC) thermograms of DPPC liposomes prepared with various compositions of 2,2’,3,3’,4,4’-HCBP (Aroclor 1260). All samples were prepared with 25 mg/mL DPPC and A) 0%, B) 1%, C) 5%, D) 10%, E) 20% by mole of the compound to DPPC. The transition temperature, T_m_, decreases with increasing HCBP content *(right)*.

### Analysis of DSC thermograms shows that 2,2’,3,3’,4,4’-HCBP destabilizes liposomes

The DSC raw tabulated data files were imported and processed using KaleidaGraph version 4.5 software. Data were normalized to zero and the baseline was subtracted. Figure 3 summarizes the DSC thermograms for each of the five samples including 0% HCBP (pure DPPC liposomes).

Analysis of the melting curves (thermograms) showed that pure DPPC liposomes had a major phase transition at 40.8 °C, which is consistent with what we expected based on previous work.^52–55^ In the presence of HCBP, however, a distinct broadening in the melting curves occurred and the peak morphology changed becoming more broad and depressed with a more pronounced peak at the T_m_.

The thermal stability of passively incorporated HCBP was also analyzed; however, there were no appreciable differences compared to pure DPPC liposomes that could be attributed to the presence of HCBP.

### Verification of 2,2’,3,3’,4,4’-HCBP incorporation into liposomes using UV-Vis analysis

To evaluate the extent of HCBP incorporation into DPPC liposomes we used UV-Vis absorption spectroscopy. There is little reported on the absorption properties of HCBP (Aroclor 1260), therefore, initially a full spectrum scan was required to establish lambda max, λ_max_. The scan showed that HCBP had a maximum absorbance at 236 nm in 200 proof ethanol. All subsequent samples were measured for HCBP incorporation at this wavelength. A standard curve was generated from which an extinction coefficient could be determined. Using the slope of the line generated in Figure 5 an extinction coefficient of 26,455 M^-1^cm^-1^ was established.^56^ To our knowledge a precise value for the extinction coefficient of Aroclor 1260 in ethanol has not been reported. We chose to investigate both the direct and passive incorporation of HCBP. Table 3 summarizes the results from both studies. After 4 hours of exposure to HCBP, the minimum exposure time, detectable levels of HCBP had already begun to appear in the liposome mixture. Despite the clear presence of HCBP in these samples it was not enough to lower the phase transition temperature and we found no appreciable changes in T_m_ from the DSC analysis (not shown). Incorporation of HCBP increased proportionally with exposure time from 4 to 24 hours, but then gradually levels off as it approaches 24 hours (Table 3). Figure 4 shows the % incorporation of HCBP in the passively absorbed samples, which never reached more than 23% by weight beginning from a 1 mg dried film. For the direct incorporation, up to 83.7% relative to the predicted theoretical quantity was determined for the 1% HCBP sample. The % incorporation decreased with increasing HCBP concentration. In all cases, the passive absorption did not significantly alter the transition temperature in the DSC for any of the samples.

**Table 3.** Quantitative analysis of 2,2’,3,3’,4,4’-HCBP incorporation into liposomes.

| Direct Incorporation |  |  |  | Passive Absorption |  |  |  |
| --- | --- | --- | --- | --- | --- | --- | --- |
| % mol DPPC | HCBP mg/mL | Theoretical HCBP, mg/mL | % Incorporated (from Theoretical) | Time, hrs | HCBP mg/mL | Residual HCBP mg/mL | % Incorporated (from 1 mg film) |
| 0 | ----- | ----- | ----- | 24 hrs – Buffer only | ~ 0.00 | ----- | ----- |
| 1 | 0.105 | 0.123 | 83.7 | 4 | 0.0241 | ----- | 2.41 |
| 5 | 0.378 | 0.615 | 61.5 | 8 | 0.130 | 0.730 | 13.0 |
| 10 | 0.621 | 1.230 | 50.5 | 12 | 0.219 | 0.645 | 21.9 |
| 20 | 0.705 | 2.460 | 28.7 | 24 | 0.229 | 0.593 | 22.9 |

\*All liposome samples were prepared with 25 mg/mL DPPC. The quantities were determined from concentrations based on absorption measurements and calculated using the extinction coefficient of 2,2,3,3',4,4'-hexachlorobiphenyl, which was found to be 26,455 $M^{-1}cm^{-1}$ from the standard curve in ethanol. Dissolution of HCBP (Aroclor 1260) into HEPES buffer was negligible after 24 hours of passive exposure.

**Figure 4.**
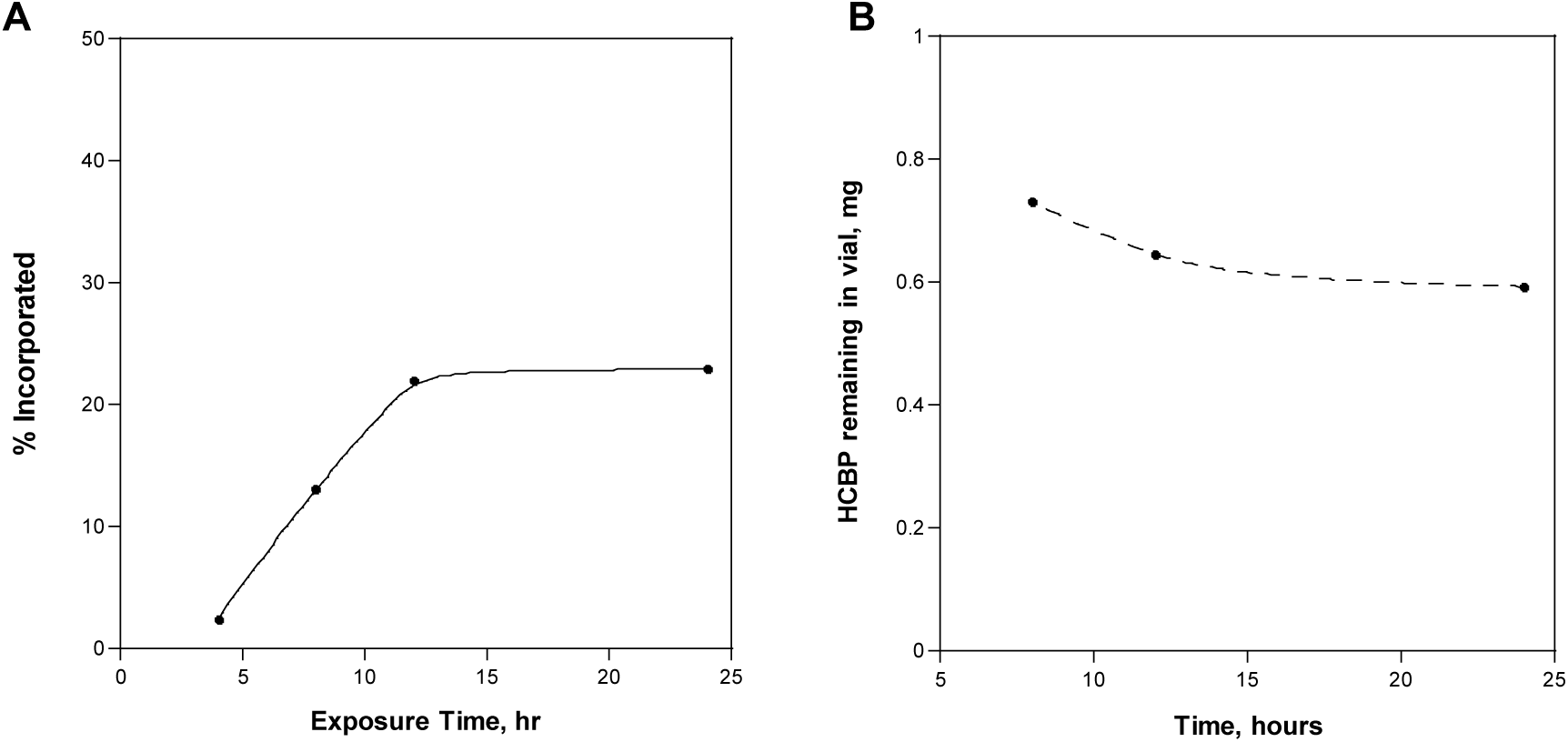
A) UV absorbance analysis of the passive incorporation of HCBP after 4, 8, 12, and 24 hours of exposure to 1 mg dried film. Incorporation increases proportionally and begins to level off after approximately 12 hours of passive exposure. B) HCBP remaining in vials after passive exposure to pure DPPC liposomes for 8, 12 and 24 hours. The amount of HCBP remaining from a dried 1 mg film decreases over time and along with the amount incorporated into liposomes represents the approximate total amount of HCBP available.

**Figure 5.**
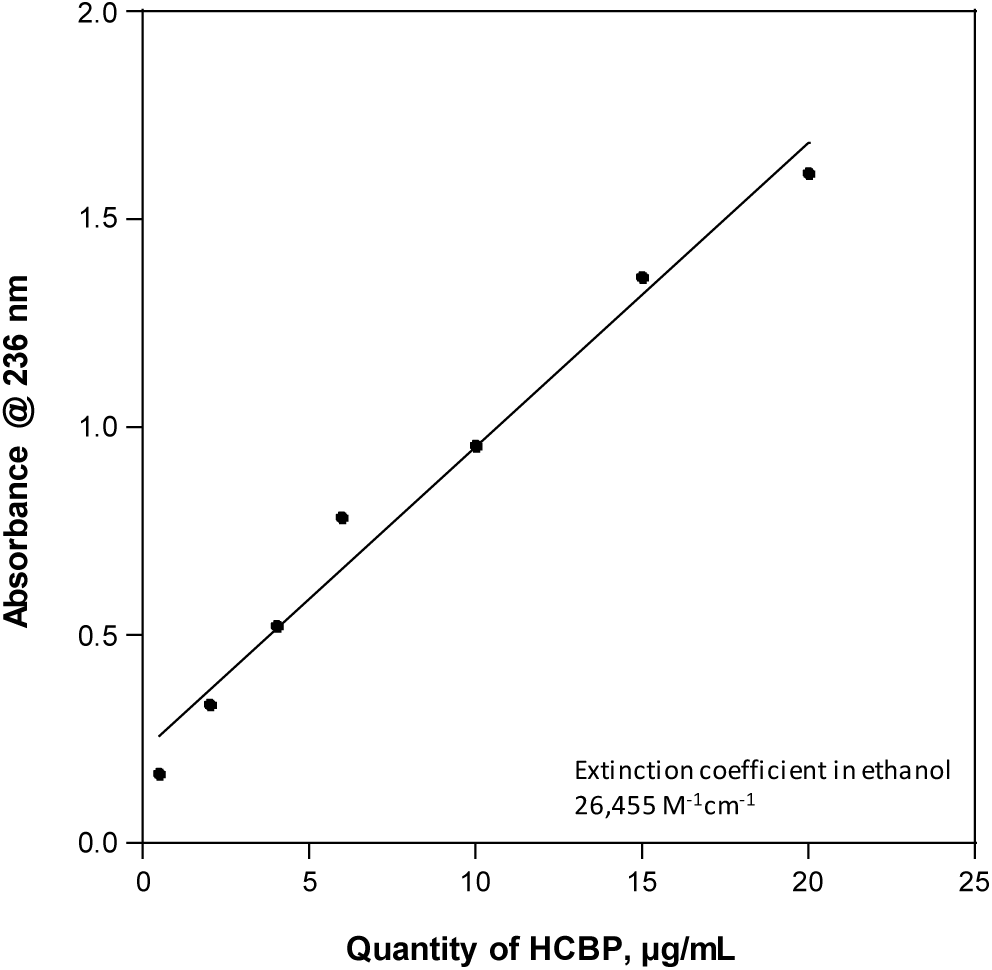
Standard curve of HCBP measured in ethanol with a calculated extinction coefficient of 26,455 M^-1^cm^-1^. The extinction coefficient was determined from the slope of the standard curve as a function of HCBP concentration (calculated from the Molarity). Subsequent analysis for HCBP levels was based on the calculated extinction coefficient applied to standard Beer’s Law.

## Discussion

We sought to investigate the effects of hexa-chlorinated biphenyl (HCBP), a congener of a class of environmental pollutants known as persistent organic pollutants on the thermal stability of DPPC liposomes using differential scanning calorimetry (DSC) and UV-Vis spectroscopy. From the DSC data we can evaluate relative liposome stability as a function of its melting temperature (T_m_) when we compare pure DPPC liposomes to those that were prepared or exposed to the environmental pollutant, HCBP. We found that when we increased HCBP content in the direct preparation the major temperature transition decreased indicating reduced thermal stability. This change was not detectable in samples that were passively exposed to HCBP, although, UV-Vis spectrophotometric analysis indicated that HCBP was present, but to a lesser degree. This is in contrast to what has been reported for compounds like cholesterol.^12,57,58^ In the presence of cholesterol a reported broadening of the major temperature transition range occurs albeit toward higher temperature.^22^ We surmise that in the absence of small molecules DPPC liposomes are free to pack more tightly with an ordering of the lipid tails giving rise to greater thermal stability. As HCBP is introduced it disrupts the lipid packing of the long chain fatty acid tails preventing them from assembling into a more ordered arrangement.^11,18,59^ Structurally, cholesterol can intercalate itself between the lipid tails of DPPC and help stabilize hydrophobic interactions in part because it is reportedly more planar and rigid.^50,60–62^ HCBP, having chlorine atoms at various substituted positions, does not have the same steric orientation that suggests it would behave in the same way. It lacks a fused ring system, which we predict gives rise to the observed thermal destabilization. In Figure 7, a schematic diagram shows how we envision and postulate HCBP incorporates into DPPC liposomes in both direct and passive preparations. The higher % incorporation from the direct preparation significantly destabilizes liposomes resulting in a lower observed T_m_. Though we did not specifically investigate how HCBP incorporation affects the size distribution, we believe it may have an effect based on the extensive peak broadening observed in the DSC thermograms (Figure 3).

**Figure 6.**
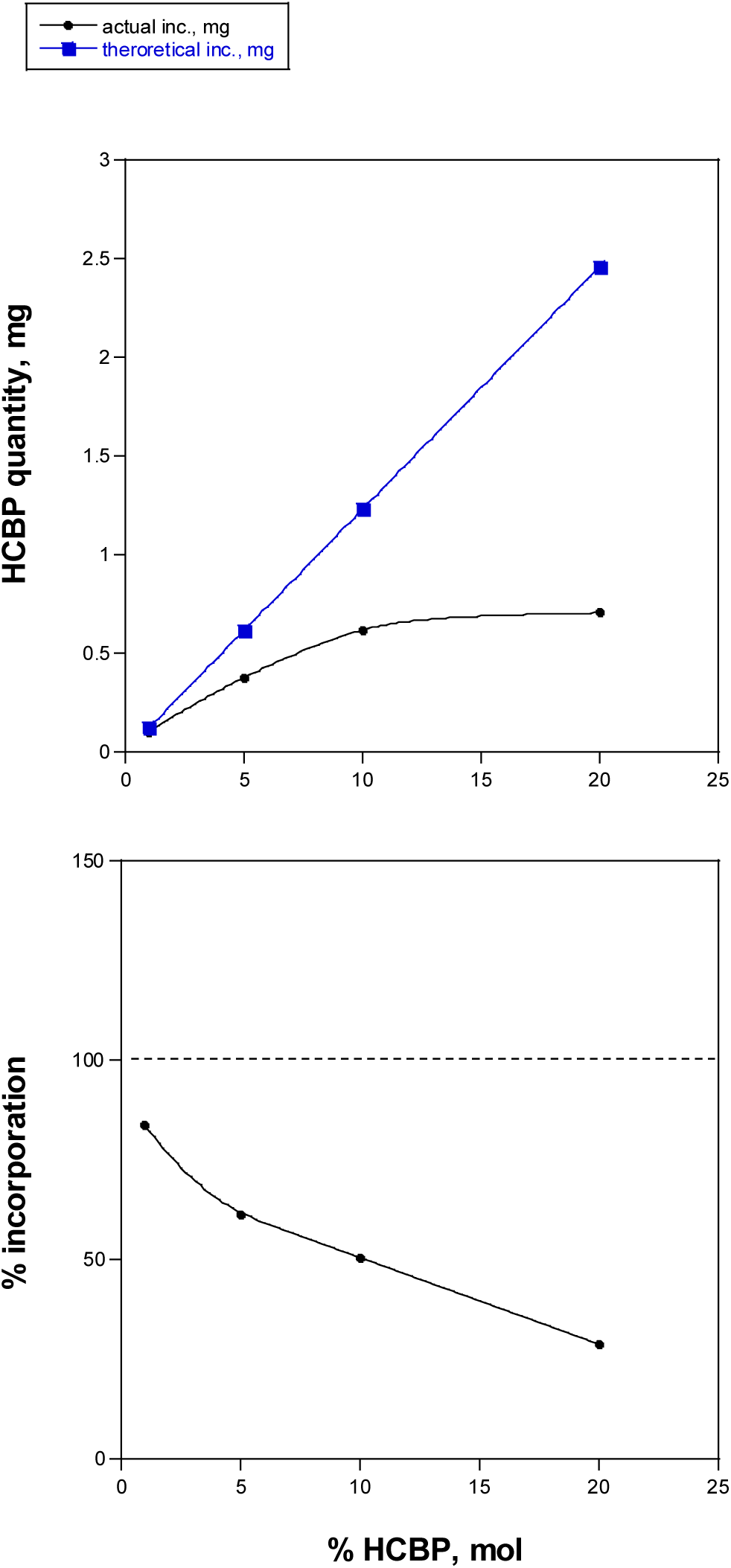
Theoretical vs. actual HCBP after direct incorporation. Experimental values were determined using the extinction coefficient generated from the standard curve and Beer’s Law was used to quantitate HCBP from the measured absorbance for each sample.

**Figure 7.**
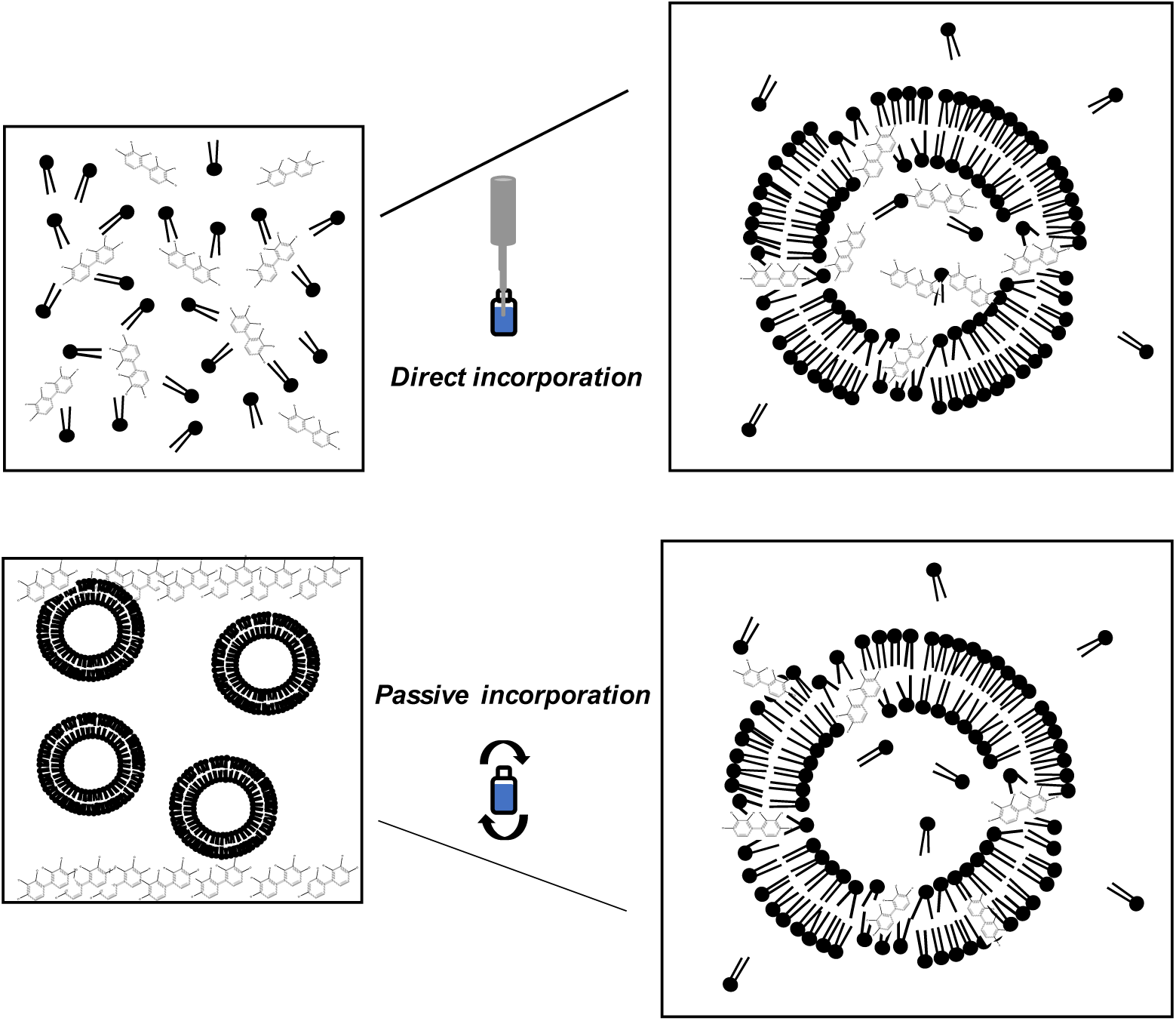
Schematic diagram of the direct and passive incorporation of HCBP into DPPC liposomes. Compiled UV-Vis data from the passive absorption suggests HCBP may preferentially partition into the lipid bilayer and not the aqueous environment based on the negligible absorption measurement in the buffer control (Table 3).

We have shown that emerging toxic environmental compounds belonging to a class of persistent organic pollutants can be incorporated into DPPC liposomes both directly and passively using a 2,2’,3,3’,4,4’-HCBP polychlorinated biphenyl compound as a representative example. The direct preparation of liposomes in the presence of this compound results in an increased loading capacity overall compared to the passive absorption method. From a practical perspective, a passive approach may be more useful in downstream applications because there are fewer technical challenges. However, the loading capacity of these liposomes is substantially lower compared to the direct method of incorporation (Table 3). Passive incorporation is less disruptive to the thermal stability overall making them more robust and potentially adaptable to a biotechnology platform. The direct incorporation of HCBP into liposomes reaches a threshold at 10% HCBP before leveling off (Figure 6). Passive incorporation shows that after 12 hours of HCBP exposure the extent of incorporation begins to diminish leaving residual behind on the vial. This is a quantitative process and it does not appear that a significant portion of HCBP leaches into the buffer itself, which could imply that there is a preference for the compound to partition into the hydrophobic bilayer of the liposome (Figure 4). Additional factors like introducing unsaturated lipids and lipids with shorter chain lengths may help to increase the compound loading capacity, which is an interesting direction to pursue.

## Acknowledgements

We would like to acknowledge the Department of Chemistry for their support.

